# Effectiveness of Exercise Intervention in Preventing Active Arthritis Exacerbation in an SKG Mouse Model of Rheumatoid Arthritis

**DOI:** 10.1101/2024.08.25.609287

**Authors:** Kaichi Ozone, Tatsunori Kumagai, Kouhei Arakawa, Takehito Sugasawa, Wenchao Gu, Sora Kawabata, Naoki Shimada, Haruna Takahashi, Moe Yoneno, Yuki Minegishi, Kei Takahata, Michiaki Sato, Yuichiro Oka, Naohiko Kanemura

## Abstract

**Objectives:** To investigate the effects of low-intensity exercise on active arthritis in an SKG mouse model of human rheumatoid arthritis (RA) pathology.

**Methods:** Twenty-four female SKG mice were divided into three groups: ‘sedentary’ (control), ‘AR’ (induced arthritis), and ‘AREx’ (induced arthritis plus low-intensity exercise). Arthritis was induced via intraperitoneal administration of mannan. After a 2-week inflammation period, low-intensity treadmill exercise was performed only in the AREx group. Arthritis was assessed weekly during the rearing period. After 4 weeks of exercise, histological and bone morphometric analyses of the right ankle joint were performed. A histological analysis of the gastrocnemius muscle was also performed. Bulk mRNA sequencing was conducted on the left synovial membrane-fat pad (SM-FP) complex.

**Results:** The synovitis score showed no change; however, the arthritis score was significantly lower in the AREx group than in the AR group (*p*<0.05), indicating that low-intensity exercise suppressed arthritis exacerbation. The calcaneal and talar bone volumes decreased in the AR group, whereas the AREx group showed no significant change. In the SM-FP complex tissue, the gene expression of inflammatory cytokines decreased in the AREx group compared with the AR group, particularly the suppression of IL6/Jak/Stat3. Immunohistochemical analysis revealed significantly decreased expression of pro-inflammatory cytokines and increased expression of anti-inflammatory cytokines in the synovium of the AREx group compared with the AR group (*p*<0.05).

**Conclusion:** Low-intensity exercise therapy for active RA showed anti-inflammatory and suppressive effects on arthritis exacerbation in SKG mice, a mouse model of human RA pathology.

**Key messages:**

- Exercise had an anti-inflammatory effect on SKG mice, a mouse model of rheumatoid arthritis.
- Exercise suppresses pro-inflammatory cytokine pathways such as IL6/Jak/Stat3 signalling in the synovial-fat complex tissue.
- Exercise therapy is effective in improving the pathophysiology of active rheumatoid arthritis.

## Introduction

Rheumatoid arthritis (RA) is an autoimmune disease, with synovitis being its hallmark feature. RA is characterized by destructive arthritis, leading to painful and irreversible deformities that drastically reduce the quality of life of affected individuals. RA has been recognized as an intractable disease since ancient times. Since the 20th century, with the advent of novel molecular therapeutics, RA has been redefined as a disease with the potential for remission. Therefore, the first treatment strategy is pharmacotherapy, with rehabilitation being additionally prescribed. However, the high cost and side effects of molecular biological drugs have led to numerous limitations in their prescription[1,2].

Recently, exercise therapy, an inexpensive non-pharmacological rehabilitation strategy, has been attracting attention as an effective means of reducing patient subjective evaluation. Human intervention studies have demonstrated the effects of exercise therapy in patients with RA. A randomized controlled trial by Durcan et al. reported improvements in patient subjective evaluation after a 12-week home exercise intervention[3]. Nevertheless, only a few studies have examined the effects of exercise on active RA. Häkkinen et al. showed that 2-year home training resulted in improved muscle strength and reduced pain among patients with early RA within 5 years of onset, and the effects lasted for 3 years[4]. Hurkmans et al. reported that dynamic exercise programs had no detrimental impact on RA; they recommended strength training and aerobic training as routine practices for patients with RA[5]. Such studies have validated the usefulness of exercise therapy as a non-pharmacological treatment for RA that does not adversely affect the disease state. However, the mechanism through which exercise has a positive effect on RA has not been studied in humans, and only a few studies have investigated the anti-inflammatory effects of exercise from histopathological and molecular biological perspectives. Preclinical validation in small animals is a prerequisite to ensure the usefulness of exercise therapy for RA.

SKG mice carrying a point mutation in the T-cell signalling molecule ZAP-70 spontaneously develop chronic arthritis under conventional microbial conditions and after the injection of a product containing β-glucan under specific pathogen-free conditions[6–8]. In contrast to collagen-induced arthritis, SKG mice not only develop arthritis but also exhibit extra-articular manifestations, such as pneumonia and rheumatoid nodules, and are a pathogenesis model that closely resembles human RA pathology, wherein Th17-mediated autoimmune arthritis spontaneously develops[6,7]. Histopathologically, the disease is characterized by synovial membrane proliferation and inflammatory cell infiltration into the subsynovial tissue, which progresses to periarthritis, pannus formation, and destruction of the cartilage and subchondral tissue[6]. SKG mice are characterized by abundant production of pro-inflammatory cytokines, such as tumor necrosis factor-alpha (TNFα), interleukin-1 beta (IL1β), and interleukin-6 (IL6) in the synovial membrane tissue[6,9], making them an appropriate laboratory animal for mimicking human RA pathology.

This study aimed to investigate the effects of exercise on RA pathology, especially in the active phase, from histopathological and molecular biological perspectives using the SKG mouse.

## Methods

This study was conducted in accordance with the University Animal Experiment Guidelines and was approved by the Animal Research Ethics Committee of our university (approval number: 2023-3).

### Experimental design and exercise intervention protocol

Twenty-four 6-week-old female SKG/Jcl mice were purchased from CLEA Japan (Tokyo, Japan). Mice were housed in cages (2 mice per cage) under a 12-hour light/dark cycle at 23 ± 1 °C. After 4 weeks of normal rearing, the mice were randomly divided into three groups at 10 weeks of age: ‘sedentary’, induced arthritis (‘AR’), and induced arthritis plus exercise (‘AREx’). For the AR and AREx groups, mannan from *Saccharomyces cerevisiae* (M-7504; Sigma-Aldrich, St. Louis, MO, USA) was dissolved in 100-mg/mL phosphate-buffered saline (PBS), and 20 mg per mouse was administered intraperitoneally to induce arthritis. The sedentary group intraperitoneally received the same volume of PBS. A 2-week normal rearing period was established as the onset period. After the onset of ear reddening and mild joint swelling, the AREx group was subjected to forced running exercise on a treadmill for small animals (Figure 1A). The exercise intensity was set at 12 m/min for 30 min/day, 3 times a week for 4 weeks to set a low intensity. Body weight, volume of unconsumed food, and arthritis scores were determined weekly during the rearing period. At 16 weeks of age, the handgrip strength of each mouse was measured using a grip strength measurement device for small animals. Arthritis scores were evaluated according to the CLEA Japan method. Details are described in the Supplementary Methods.

**Figure 1.**
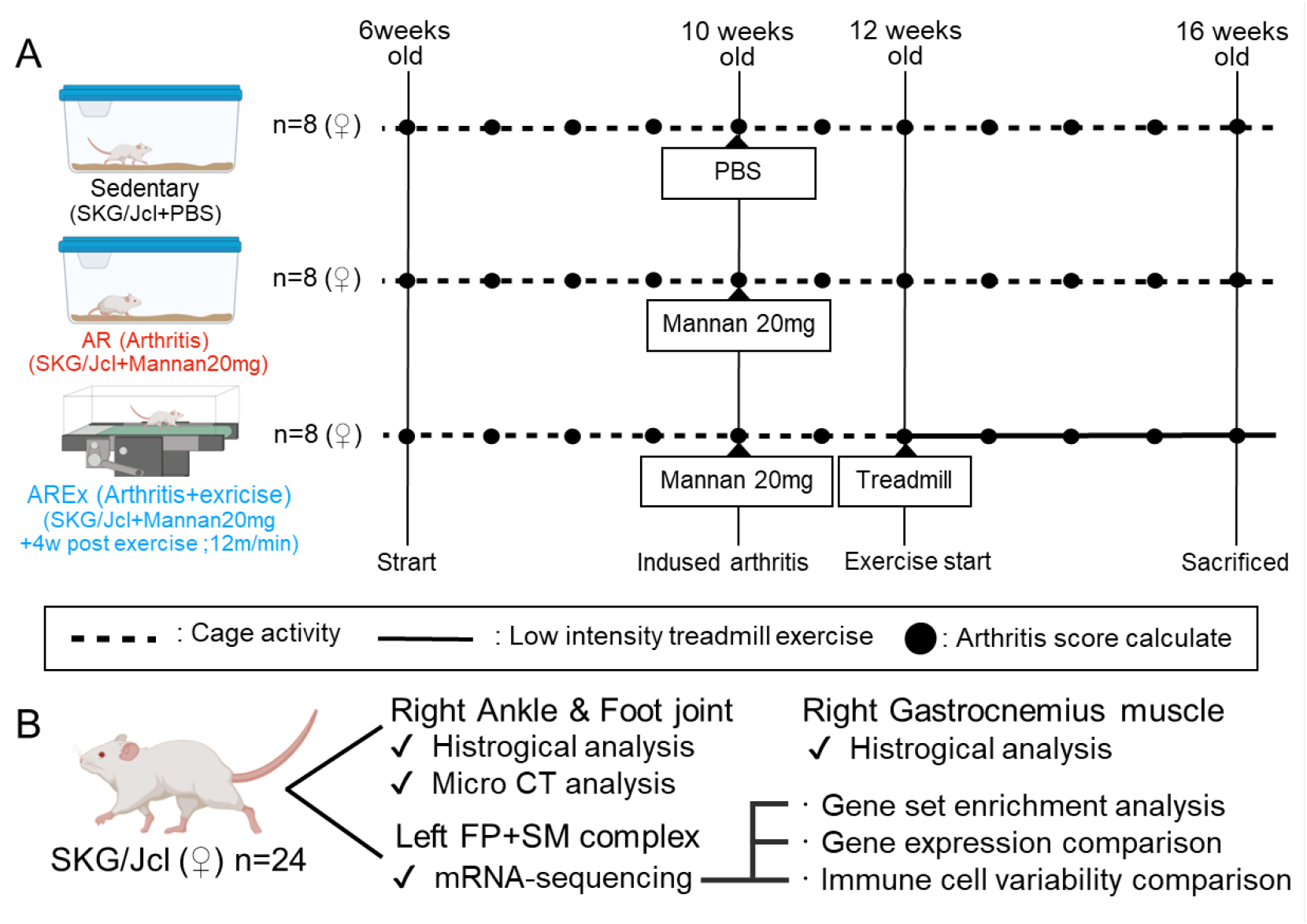
Study design and tissue collection. (A) SKG mice were divided into three groups: the sedentary group was a control group that received phosphate-buffered saline (PBS); the AR group was induced with arthritis and allowed to free range; and the AREx group was induced with arthritis followed by 4 weeks of low-intensity exercise. All mice were free for four weeks, after which only the AR and AREx groups developed arthritis. Arthritis was induced by intraperitoneal administration of 20 mg mannan. After a 2-week arthritis-induction period, only the AREx group underwent low-impact exercise on a treadmill. Dotted line, free ranging; solid line, treadmill exercise; black circles, time of arthritis score measurement. (B) Tissue and samples collected from 24 SKG mice. Created with BioRender.com.

After each measurement, the mice were humanely euthanized via cervical dislocation under deep anaesthesia using isoflurane. Immediately after euthanasia, the right ankle joint was harvested for morphological and histological analyses. The right gastrocnemius muscle was also harvested for histological analysis. The synovial membrane-fat pad (SM-FP) complex tissue was harvested from the left ankle joint under a stereomicroscope for molecular biology analysis (Figure 1B).

### Morphological analysis

Ankle joints were fixed in 4% paraformaldehyde solution for 24 h. Micro-computed tomography (CT) was performed on representative ankle joints (n=15) of the total samples using SKYSCAN 1272 (Bruker, Billerica, MA, USA). The measurement conditions for the X-ray, detector resolution, pixel size, rotation angle pitch, and filter setting were 60 kVp/165 μA, 1632 × 1092, 16.2 μm, 0.4 deg/s, and 0.5 mm aluminium. The measured data were converted to 3D data using a dedicated visualization application (CTVox; Bruker) and analyzed using the CT Analyzer application (Bruker) [10,11]. On sagittal cross-sectional images, the bone volumes (BV) of the calcaneus and talus were calculated by setting a region of interest with a free pen.

### Histological analysis

The left gastrocnemius muscle was collected (n=24), infiltrated with isopentane, and then frozen at −100 °C. Subsequently, the muscle was sliced at 12 μm. Haematoxylin and eosin (HE) staining was performed on frozen sections to compare the muscle cross-sectional area (CSA). The CSA was measured using Fiji image analysis software[12], and the average muscle CSA per muscle fibre was calculated.

After bone morphometric analysis, the right ankle joint was infiltrated with 14% ethylenediaminetetraacetic acid solution and demineralized for 2 weeks (n=24). Subsequently, they were paraffin-replaced and thinned to a thickness of 5 μm. After deparaffinization, HE staining and toluidine blue (TB) staining were performed. Tartrate-resistant acid phosphatase (TRAP) staining was also performed using a TRAP/alkaline phosphatase staining kit (FUJIFILM Wako Pure Chemical Co., Osaka, Japan) according to the manufacturer’s instructions. The HE-stained right ankle joint was blinded to the samples for synovitis assessment, and three proficient raters calculated the synovitis scores. The synovitis score is scored 0–3 points for each of the following three items, with a total score of 0–9[13]: enlargement of the synovial lining cell layer, density of the resident cells, and inflammatory infiltrate. Articular cartilage staining was evaluated in TB-stained tissues. The tibiotalar and calcaneocuboid joints were evaluated, the percentage of the TB-positive area in the articular cartilage was measured using Fiji image analysis software[12], and the number of osteoclasts was counted in the TRAP-stained sections. To calculate the area of bone erosion, regions of interest were defined as the areas of synovial membrane infiltration above the calcaneus and the cubital joint of the calcaneus. Osteoclast counts and bone erosion area calculations were performed using the Fiji image analysis software[12].

### Bulk mRNA sequencing in the SM-FP complex

The SM-FP complex tissue from the left ankle joint was immediately infiltrated into RNAlater™ Stabilization Solution (Invitrogen, Waltham, MA, USA) and stored at −20 °C (n=24). At the time of analysis, tissues from two individuals were pooled into one group, with four pools per group. The tissues were infiltrated with QIAzol Lysis Reagent (QIAGEN, Hilden, Germany), pulverized with beads, and total RNA was extracted using the miRNeasy Micro kit (QIAGEN). Subsequently, mRNA-sequence analysis was requested by the Open Facility Promotion Organization of the University of Tsukuba, and FASTQ files were obtained. Details of the analysis are described in the Supplementary methods.

### Bioinformatic analysis

Bioinformatic analysis was performed using CLC Genomics Workbench software to extract the genes of interest. Volcano plots were generated to detect significant differences in gene expression between the AR and AREx groups. Genes of interest were profiled using the RNA-seq data controller, highlighting expression features (RNA-seq chef)[14], a web-based integrated transcriptome analysis platform. The trimmed mean M values (TMM) and transcripts per million (TPM) were normalized to the total read counts of the extracted genes of interest in the AR and AREx groups. The TMM-normalized values were used for gene set enrichment analysis (GSEA). GSEA was performed by selecting the Hallmark gene set from the Mouse Collection (MSigDB) and setting the AR group as ‘Class A’ and the AREx group as ‘Class B’. The gene set database used in the degree of enrichment was quantified as an enrichment score (ES). A normalized ES (NES) was calculated by normalizing the ES according to the size of the gene set. For the comparison of gene expression, TPM-normalized values were used between groups. The mouse immune cells were calculated by the single-sample GSEA (ssGSEA) method using a gene set based on the R package of ‘ImmuCellAI-mouse’[15,16] Details of the analysis are described in the Supplementary Text.

### Immunohistochemical (IHC) analysis

For IHC staining, anti-CD4, anti-TNF-α, anti-IL6, anti-IL1β, anti-IL4, and anti-IL10 were used as primary antibodies. For details, see the Supplementary Materials and Methods. The area ratio of positive regions per unit area of IHC-stained images was calculated using the Hybrid Cell Count Application (BZ-H4C, KEYENCE, Osaka, Japan) in the BZ-X Analyzer software (BZ-H4A, KEYENCE).

### Statistical analysis

Statistical analysis was performed using JASP 0.16.4[17]. After determining the normality of the distribution of each dataset, a one-way analysis of variance (ANOVA) was performed to confirm the effect of the intervention. Two-way ANOVA was applied to the RA score, body weight, and volume of unconsumed food. The Kruskal–Wallis test was performed only to compare synovitis scores. The Bonferroni method was used for all post-hoc tests. Parametric data were presented as mean ± standard deviation, whereas non-parametric data were expressed as medians (interquartile ranges). Statistical significance was set at *p*<0.05.

## Results

### Exercise mildly inhibited arthritis exacerbation and prevented muscle atrophy

Macroscopic images of the hands and feet at 16 weeks of age and bone CT images of the entire foot are shown in Figure 2A. Trends in body weight and food intake are shown in Supplementary Figure 1A and 1B. None of the groups showed significant changes at 6–10 weeks of age, confirming that the disease did not develop spontaneously. From the 11-week post-injection time-point, arthritis was significantly induced in the AR and AREx groups compared to that in the sedentary group (*p<*0.05; Figure 2B). At 13 weeks after the start of the exercise intervention, there was already a significant difference between the AR and AREx groups, and this trend continued until 16 weeks of age (*p<*0.05; Figure 2B). At 16 weeks, the talus bone volume significantly decreased in the AR group compared with that in the sedentary group (*p<*0.05; Figure 2C). Similarly, the calcaneus bone volume significantly decreased in the AR group compared to that in the sedentary group (*p<*0.05; Figure 2D).

**Figure 2.**
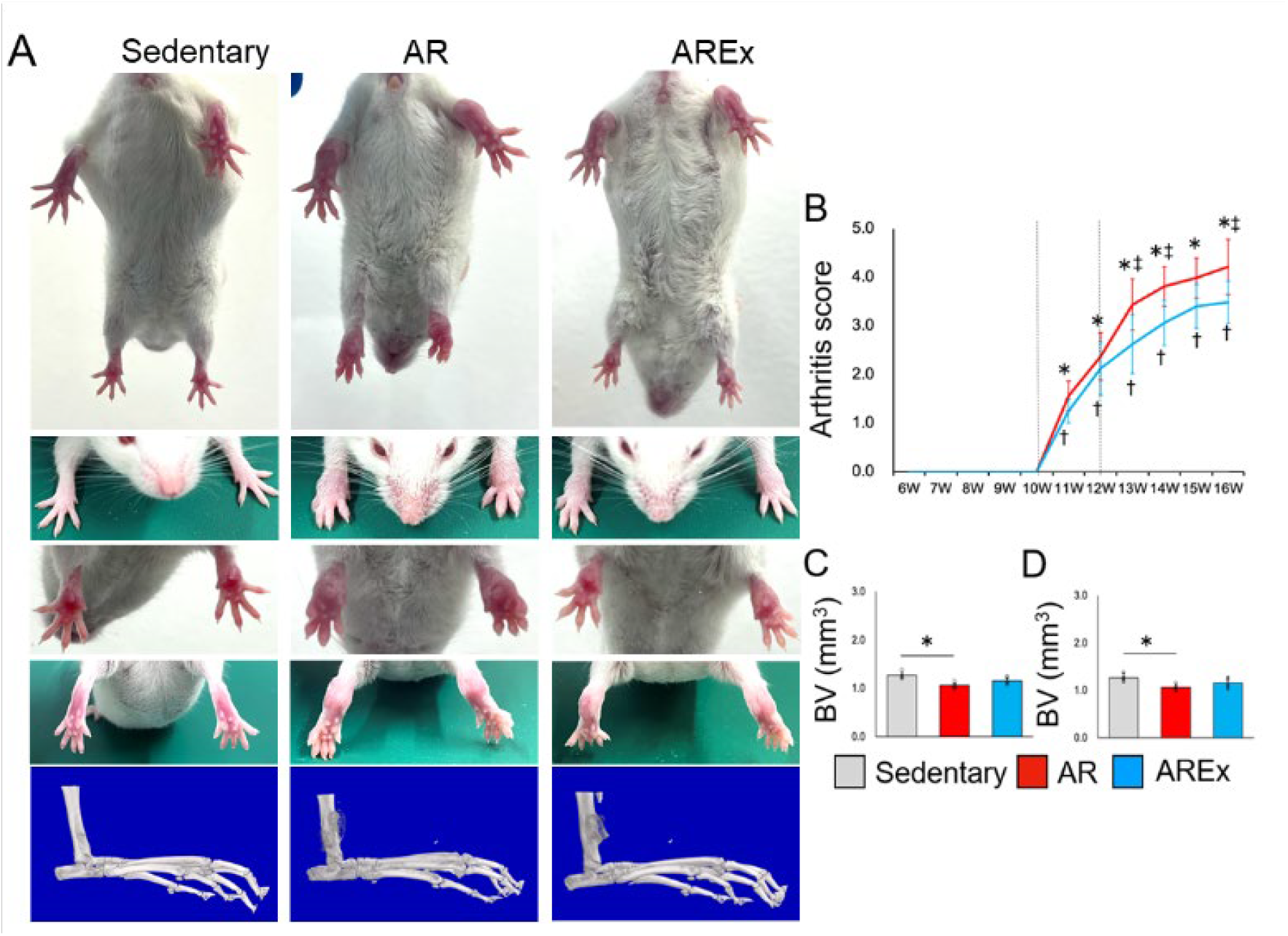
Arthritis variation and bone morphology analysis in SKG mice. (A) Comparative arthritis images of the wrist and ankle joints and ankle joint images obtained using micro-CT are shown. (B) Variation in the arthritis score for each individual calculated weekly. Results of bone volume calculations for the talus (C) and calcaneus (D). Grey, red, and blue indicate sedentary, AR, and AREx groups, respectively. *; sedentary vs AR: *p*<0.05, †; sedentary vs AREx: *p*<0.05, ‡; AR vs AREx: *p*<0.05.

The degree of arthritis prevention and exercise-related muscle atrophy was evaluated by calculating the cross-sectional muscle area of the gastrocnemius muscle and the handgrip strength, which reflects total body muscle strength (Supplementary Figure 2A). The mean CSA of the gastrocnemius muscle normalized by body weight was significantly reduced in the AR group, with significant differences between the sedentary and AR groups and between the AR and AREx groups (*p<*0.05). The AREx group had the same area as the sedentary group, and no significant difference was observed between the AREx and sedentary groups (*p*>0.05; Supplementary Figure 2B). Similar results were obtained for the handgrip strength (Supplementary Figure 2C).

### Exercise did not improve synovitis but protected the articular cartilage

HE-stained images of the entire ankle joint area, TB-stained images of the tibiotalar bone joint, and TB-stained images of the calcaneocuboid joint are shown in Figure 3A–C. Synovial membrane thickening, cell proliferation, and inflammatory cell infiltration were observed in both the AR and AREx groups. The synovitis scores were 0.0 (0.0–0.08) for the sedentary group, 7.62 (6.92–8.08) for the AR group, and 6.50 (5.25–7.08) for the AREx group; the AR and AREx groups scored significantly higher than the sedentary group (*p<*0.05), and no significant changes were observed between the AR and AREx groups (Figure 3D). The percentage of TB-stained area in the cartilage region was calculated to confirm the extent of proteoglycan loss in the articular cartilage. In the tibiotalar joint, significant differences were observed among all groups (Figure 3B and 3E). In the AR group, the TB-stained areas were reduced in the shallower layer than in the tidemark on both sides of the tibia and talus, whereas in the AREx group, more stained areas remained on the talar side. Similarly, significant differences in the calcaneocuboid joint were observed among all groups (Figure 3C and 3F). In the AR group, TB staining was sparsely reduced in the outer portion of the shallower layer than the tidemark, whereas in the AREx group, staining in the shallower layer than the tidemark was preserved.

**Figure 3.**
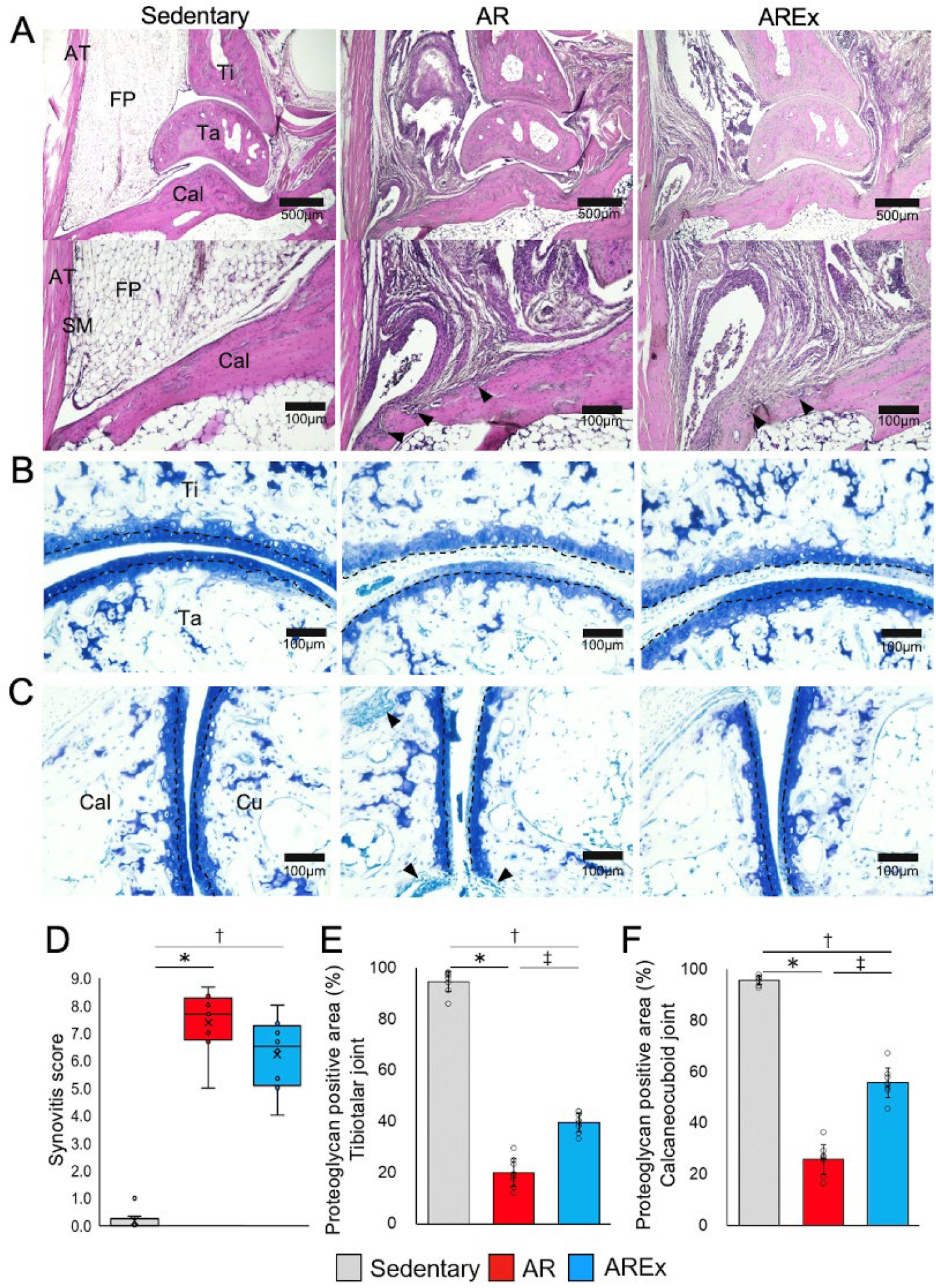
Histological evaluation of synovitis and articular cartilage. (A) The upper and lower panels show H&E-stained images of the ankle joint taken at 40× and 200× magnification, respectively. Arrowheads indicate bone erosion of the synovial membrane. AT, Achilles tendon; FP, fat pad; Ti, tibia; Ta, talus; Cal, calcaneus; SM, synovial membrane. (B) Toluidine blue-stained images of the tibiotalar joint taken at 100× magnification are shown. Ti, tibia; Ta, talus; TM, tidemark. (C) Toluidine blue-stained images of the calcaneocuboid joint taken at 100× magnification are shown. Arrowheads indicate sites of synovial membrane bone erosion. Cal, calcaneus; Cu, cuboid bone. (D) Results of the synovial membrane score calculation are shown. Three skilled raters evaluated the blinded images. (E) Results of proteoglycan-positive area ratio comparisons in the tibiotalar joint. (F) Results of proteoglycan-positive area ratio comparisons in the calcaneocuboid joint. Grey indicates the Sedentary group, red indicates the AR group, and blue indicates the AREx group. *; Sedentary vs AR: *p*<0.05, †; Sedentary vs AREx: *p*<0.05, ‡; AR vs AREx: *p*<0.05.

### Exercise suppressed osteoclast activation in the synovial membrane and prevented the worsening of bone erosion

TRAP-stained images of the calcaneal (superior) synovial and calcaneocuboid joints are shown in Figure 4A and 4B. The activation of synovial membrane osteoclasts in the calcaneal-synovial membrane space was particularly pronounced in the AR group, and bone infiltration of the synovial membrane and areas of bone erosion increased. The activated synovial membrane osteoclasts and bone erosion area were significantly less in the AREx group than in the AR group. However, bone invasion of the synovial membrane was also observed (*p*<0.05; Figure 4A and 4C). Similar results were observed in the calcaneocuboid joint (Figures 4B and 4D).

**Figure 4.**
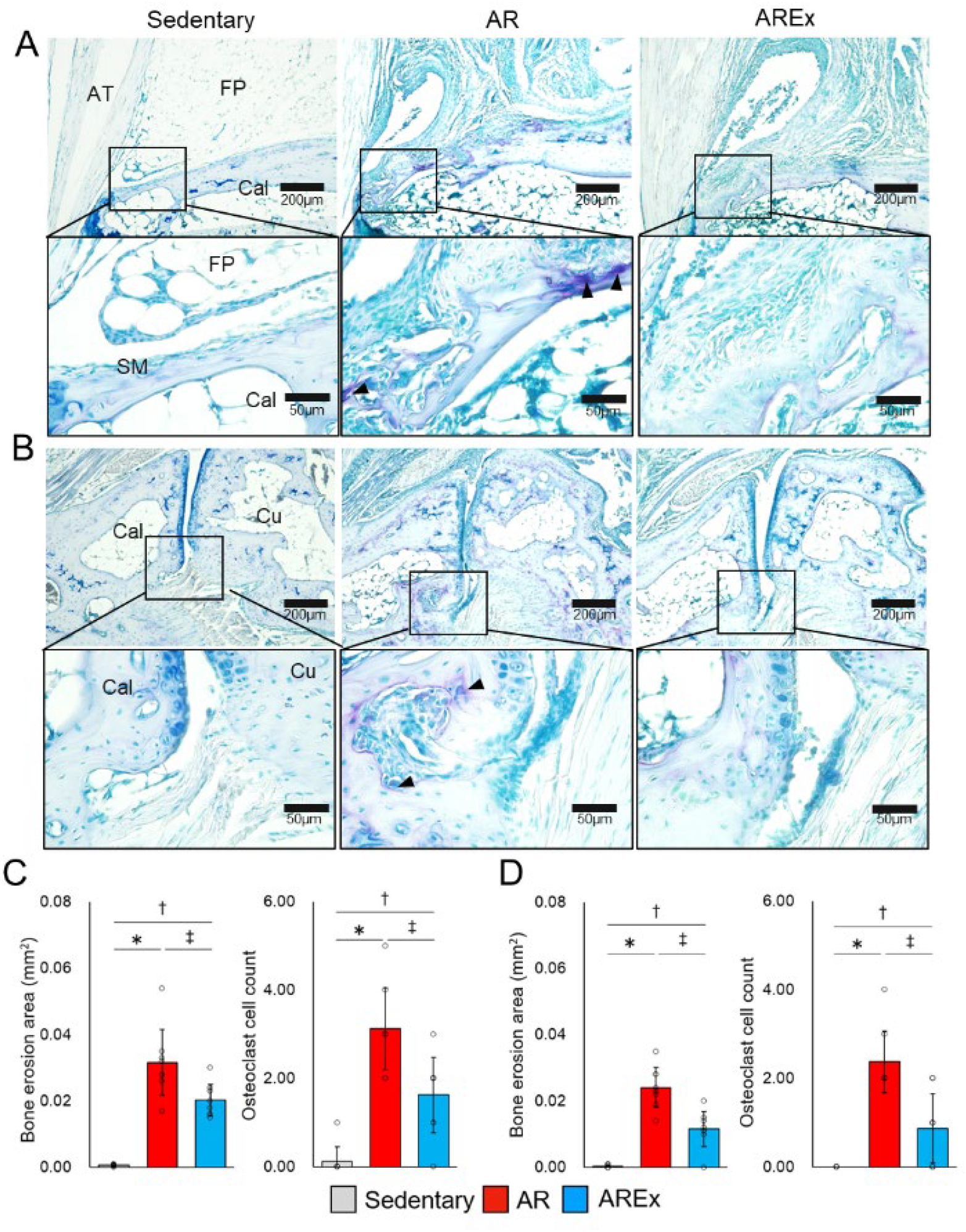
Analysis of osteoclast activation and areas of bone erosion. (A) The upper and lower panels show TRAP-stained images of the ankle joint taken at 100× and 400× magnification, respectively. Arrowheads indicate osteoclasts. AT, Achilles tendon; FP, fat pad; Cal, calcaneus. (B) TRAP-stained images of the calcaneocuboid joint taken at 100× magnification in the upper panel and 400× in the lower panel. Arrowheads indicate osteoclasts. Cal, calcaneus; Cu, cuboid bone. (C) TRAP-stained images of the ankle joint showing areas of bone erosion and osteoclast count. (D) TRAP-stained image of a calcaneocuboid joint showing areas of bone erosion and osteoclast count. Grey indicates the Sedentary group, red indicates the AR group, and blue indicates the AREx group. *; Sedentary vs AR: *p*<0.05, †; Sedentary vs AREx: *p*<0.05, ‡; AR vs AREx: *p*<0.05.

### Exercise suppressed the activation of inflammatory cytokines in the SM-FP complex tissue

Bulk mRNA sequences were obtained from the SM-FP complex tissue of the left ankle joint. Compared with the sedentary group, the AR and AREx groups showed different genetic variability. Some samples in the AREx group showed similar trends to those in the AR group, whereas others behaved differently (Figure 5). A comparison of gene expression between the AR and AREx groups confirmed a significant decrease in factors related to cytokines, such as *Ccl20, Il1r1*, and *Cxcl2* (Figure 5A). GSEA of the AR and AREx groups was performed using TMM-normalized values of raw bulk mRNA sequence data. Comparison with Hallmark gene sets revealed variation in the following representative gene sets (Figure 5B): TNFA SIGNALING VIA NFKB (NES=2.32, FDR *q<*0.001), IL6 JAK STAT3 SIGNALING (NES=2.30, FDR *q*<0.001), and INFLAMMATORY RESPONSE (NES=2.25, FDR *q*<0.001), and HYPOXIA (NES=1.43, FDR *q*<0.001). Other representative data compared with the hallmark gene set and Gene Ontology (biological process, molecular function) are shown in Supplementary Figure 3. Relative gene expression comparison using TPM-normalized values of raw mRNA sequence data revealed that *Il6, Jak1, Stat3*, and *Tnfsf11* (*Rankl*) levels were significantly lower in the AREx group than in the AR group (*p*<0.05; Figure 5C). Other representative data comparing the relative gene expression with TPM-normalized values are shown in Supplementary Figure 4. The variability of each immune cell was also calculated from the raw bulk mRNA sequence data, but no significant variability between the AR and AREx groups was identified (Supplementary Figure 5).

**Figure 5.**
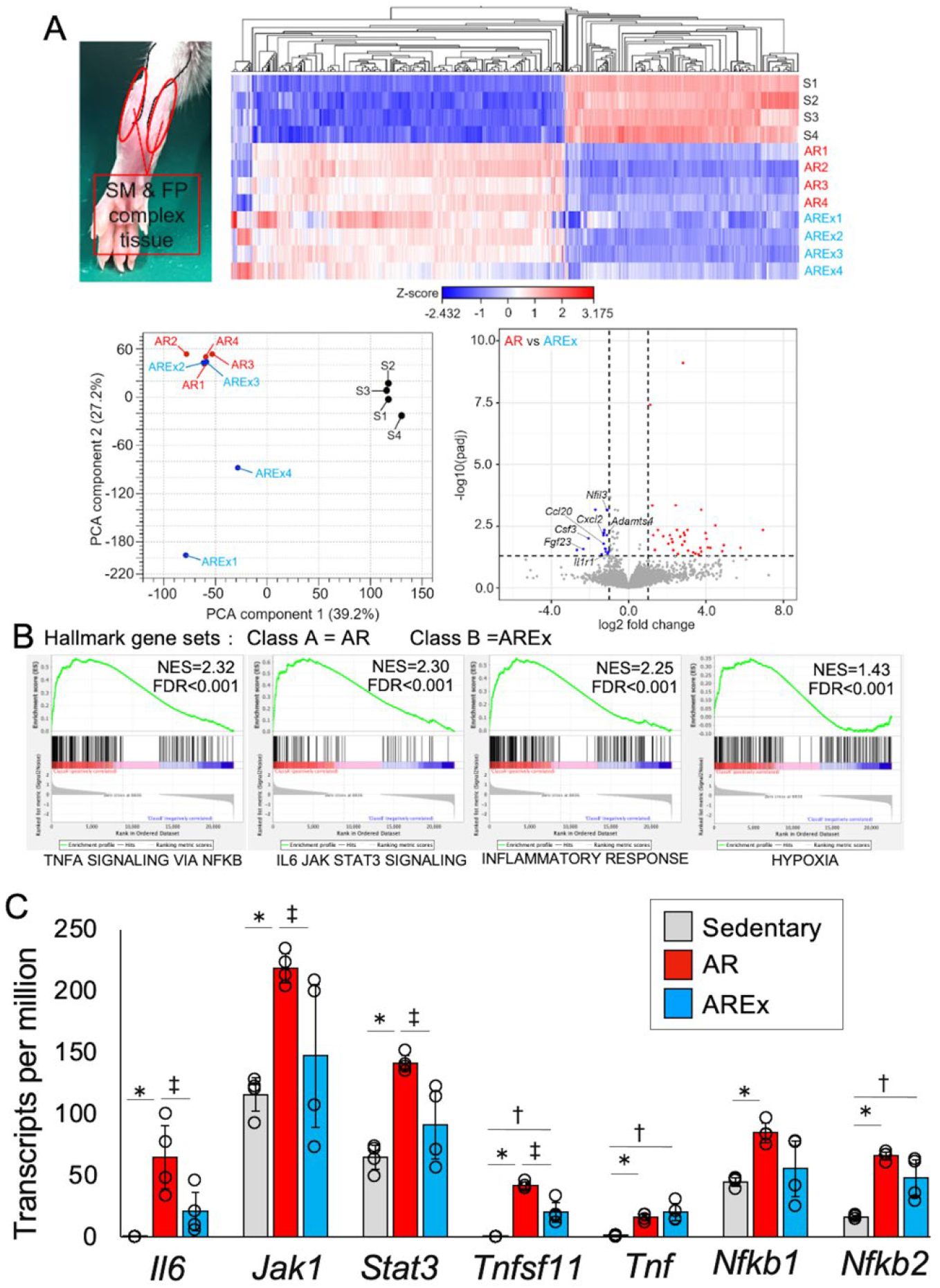
Bulk mRNA sequence results and bioinformatic analysis in the synovial membrane-fat pad (SM-FP) complex tissue. (A) Tissue samples obtained from the SM-FP complex. Tissue samples from two mice were pooled into one sample, and four samples per group were used for the analysis. Bulk mRNA sequencing was performed, and genetic variation in each sample was visualized using a heat map and principal component analysis. The AR and AREx groups were compared, and a volcano plot was constructed to identify and visualize the significantly altered genes. (B) TMM-normalized data from the total count values obtained from the bulk mRNA sequence. Hallmark gene sets ranked high in the two-group comparison between the AR and AREx groups using GSEA. Red indicates the AR group as Class A, whereas blue indicates the AREx group as Class B. Normalized enrichment score (NES) and false discovery rate (FDR) values are shown. (C) Between-group comparison of specific gene variations using TPM-normalized values obtained from the bulk mRNA sequence. Grey indicates the Sedentary group, red indicates the AR group, and blue indicates the AREx group. *; Sedentary vs AR: *p*<0.05, †; Sedentary vs AREx: *p*<0.05, ‡; AR vs AREx: *p*<0.05.

The protein expression levels of CD4-positive cells, inflammatory cytokines (TNF-α, IL6, and IL1β), and anti-inflammatory cytokines (IL4 and IL10) in the SM-FP complex tissue were compared semi-quantitatively by IHC staining. Immunohistochemical staining images are shown in Figure 6. The AREx group had a significantly lower expression of inflammatory cytokines (*p<*0.05) and a significantly higher expression of anti-inflammatory cytokines (*p<*0.05) than the AR group.

**Figure 6.**
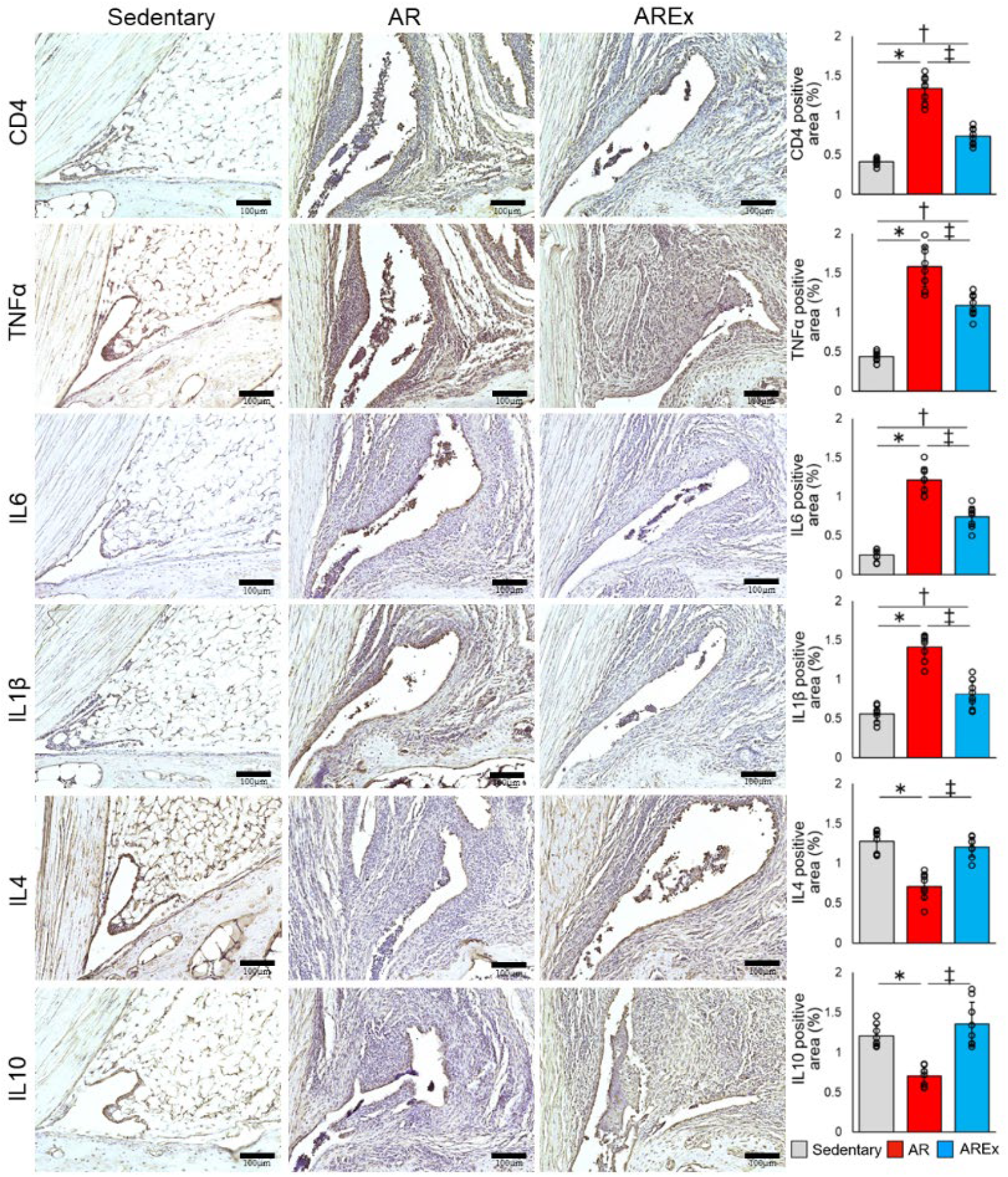
Comparison of inflammatory and anti-inflammatory cytokine expression in the synovial membrane tissue. The expression of CD4-positive T cells, inflammatory cytokines (TNF-α, IL6, IL1β), and anti-inflammatory cytokines (IL4, IL10) in the synovial membrane tissue was compared using immunohistochemical staining images. The positive area ratio of each protein in the synovial membrane was also calculated. Arrowheads indicate positive cells. Grey indicates the Sedentary group, red indicates the AR group, and blue indicates the AREx group. *; Sedentary vs AR: *p*<0.05, †; Sedentary vs AREx: *p*<0.05, ‡; AR vs AREx: *p*<0.05.

## Discussion

The present study examined the effects of exercise therapy on active RA by subjecting SKG/Jcl mice to low-intensity treadmill exercise. The results indicated that arthritis exacerbation was mildly suppressed and that mRNA and protein expression associated with inflammatory cytokines in the SM-FP complex tissue was reduced. Although exercise did not cause significant histological changes in synovitis, histological, morphological, and molecular biological perspectives indicate that exercise had a protective effect against cartilage and bone destruction.

The exercise intensity employed in this study was 12 m/min, which is a validated low-intensity (approximately 30–50% of maximal capacity) and aerobic exercise speed for rodents[18–20]. Aerobic exercise of an appropriate intensity results in mild muscle strengthening and cardiorespiratory effects[21]. Moderate treadmill running in arthritis models also ameliorates synovitis[19], and moderate mechanical stress on the articular cartilage and subchondral bone inhibits joint destruction and improves symptoms[22,23]. However, high-intensity treadmill running reportedly has several adverse effects on diseases such as osteoarthritis[24,25]. As RA is a painful disease with a high degree of joint infiltration, including inflammatory cells, intense dynamic or weight-bearing exercises are inappropriate for joint protection as they may exacerbate the disease[26]. However, several recent studies using rodent models mimicking arthritis have reported that low-intensity aerobic exercise improves the pathophysiology of arthritis[18,19,27]. In humans, exercise does not adversely affect the pathophysiology of RA, thereby increasing the significance of exercise therapy in RA[28–30]. The results of this study support the findings of previous studies and indicate that exercise is effective in preventing disease exacerbations even in active RA with increased inflammation.

This study showed that low-intensity exercise prevented muscle atrophy and mainly suppressed the activation of TNF-α signalling via NF-κB and IL6-Jak-Stat3 signalling, which are key factors in RA pathology, in the synovial membrane and adipose tissue around joints. Significant reductions in inflammatory cytokines were also observed in gene expression levels and IHC. Levels of the anti-inflammatory cytokines IL4 and IL10 were also significantly increased by low-intensity exercise at the protein level, although there were no significant changes at the gene level. The decrease in inflammatory cytokines and increase in anti-inflammatory cytokines with exercise were similar to those reported in previous studies[31–33]. Despite the greater load exerted on the joints, bone destruction and cartilage degeneration were prevented in the low-intensity exercise group. The SM-FP complex tissue, inhibition of matrix-degrading enzymes (*Mmps*), activation of anti-matrix degrading enzymes (*Timp1*), and RANKLE (*Tnfsf11*) expression were also suppressed (Supplementary Figure 4). Synovitis is the main pathogenesis of RA. Within the synovial membrane, infiltration of inflammatory cells such as lymphocytes and monocytes, marked angiogenesis, and proliferation of synovial membrane surface cells are observed[34]. Infiltrating hematopoietic cells, synovial fibroblasts, and T cells have been shown to induce the expression of various inflammatory cytokines, including MMPs, VEGF, and RANKL[35–37]. Synovial bone erosion has also been reported to be dependent on the production of RANKL (a member of the TNF family), which is essential for osteoclast formation and activity, and that inflammatory cytokines are also important for osteoclast culture[19,38]. In the present study, the histological changes in synovitis were not significant; however, the exercise group had more anti-inflammatory cytokine protein expression regions in the synovium, and inflammatory cytokine gene expression was also reduced. It is possible that synovial membrane inflammation is reduced by exercise, resulting in changes in genes associated with cartilage and bone destruction in the SM-FP complex.

A possible reason inflammatory cytokine levels are reduced by exercise is the involvement of myokines. In the present study, exercise prevented muscle atrophy; the muscle CSA increased more in the exercise group than in the sedentary group, and grip strength did not weaken. Myokines are peptide proteins produced, expressed, and released by myocytes in muscle fibres during muscle contraction, and are known to increase heat production in adipose tissue with anti-inflammatory and insulin-sensitizing effects throughout the body[39,40]. IL6, a representative myokine, is upregulated in the muscle tissue during exercise and promotes the generation of two cytokines with anti-inflammatory effects (IL1ra and IL10)[41]. In other words, exercise mediates anti-inflammatory signals that are likely to be partly mediated by IL6. Although IL6 concentrations in the muscle and plasma were not directly measured in this study, a significant increase in the anti-inflammatory cytokine IL10 protein and a decrease in inflammatory cytokines confirmed muscle hypertrophy, and activation of fatty acid metabolism was predicted, suggesting a partial myokine effect.

This study has some limitations. Tissue collection was performed at only one time-point. This study demonstrated that exercise had an inhibitory effect on inflammatory progression in active RA. However, because we did not follow the data for 4 weeks during the exercise period, we do not know how immune cells behave immediately after the start of exercise or how inflammatory and anti-inflammatory cytokines behave. Therefore, the mechanisms underlying the anti-inflammatory effects of exercise on active RA remain unelucidated. Further studies are warranted to follow the time course in the future. In addition, since the bulk mRNA sequence of the SM-FP complex tissue was analyzed in this study, tissue-specific responses and spatiotemporal expression behaviour are unknown.

Spatiotemporal transcriptome analysis will verify tissue specificity and the spatiotemporal expression of gene behaviour. In this study, we performed forced running on a treadmill to mimic low-intensity exercise in mice. However, we must carefully consider what type of low-intensity exercise is best for patients with RA since pain may be a confounding factor.

## Acknowledgements

We would like to thank Editage (www.editage.com) for English language editing.

## Data availability statement

The data underlying this article are available in the article and in its online supplementary material. In addition, raw and processed data on Bulk mRNA sequences are also stored in the Gene Expression Omnibus database (accession number GSE271058).

