## Supplemental file for "Effectiveness of Exercise Intervention in Preventing Active Arthritis Exacerbation in an SKG Mouse Model of Rheumatoid Arthritis"

### Supplementary material

#### Supplementary Data S1. Sample size

The G*Power software Version 3.1.9.6 was used to estimate the required sample size. One-way ANOVA was used to analyze histological and morphological changes after the intervention. Based on the results of a small preliminary study with the same group setting (primary outcome: RA score at 11 weeks), an F-test (ANOVA: fixed effects, omnibus, one-way ANOVA) was used. The partial η2 obtained from previous results was 0.735, so the effect size f was calculated to be 1.665408. The power was set at 80% and the alpha was set at 0.01. Based on the above information, a minimum of 12 mice was required for the three-group setup. Therefore, the sample size for each analysis was at least 4 mice.

#### Supplementary Data S2. Arthritis scores

The arthritis score was calculated as follows according to the method outlined by CLEA Japan: (i) fingers were assigned a score of 0.1 if they were red and swollen; (ii) the wrist was given a score of 0.5 if the hollow at the wrist joint was no longer present and a score of 1.0 if it protruded further; and (iii) the ankle was allocated a score of 0.5 if the Achilles tendon attachment area was obtuse and no longer indented around the Achilles tendon and a score of 1.0 if it protruded further.

#### Supplementary Data S3. Bulk mRNA sequencing methods

The SM-FP complex tissue from the left ankle joint was immediately infiltrated into RNAlater™ Stabilization Solution (Invitrogen, Waltham, MA, USA) and stored at −20 °C (n=24). At the time of analysis, tissues from two individuals were pooled into one group, with four pools per group. The tissues were infiltrated with QIAzol Lysis Reagent (QIAGEN, Hilden, Germany), pulverized with beads, and total RNA was extracted using the miRNeasy Micro kit (QIAGEN). After liquid-phase extraction, the QIAcube (QIAGEN) automatically performed the subsequent steps. The yield and absorbance of the extracted total RNA were measured using a NanoDrop One spectrophotometer (Thermo Fisher Scientific, Waltham, MA, USA). RNA integrity was checked using the Agilent RNA 6000 Nano Kit (Agilent Technologies, Santa Clara, CA, USA) on a bioanalyzer. All the RNAs were confirmed to be intact. Libraries for mRNA sequencing were created using the NEBNext Ultra II RNA

Library Prep Kit for Illumina and NEBNext Poly(A) mRNA Magnetic Isolation Module (New England Biolabs, Ipswich, MA, USA) according to the manufacturer's instructions. The concentration and size distribution of the libraries were measured using an Agilent DNA 7500 kit (Agilent Technologies) on a bioanalyzer. All samples were subjected to NGS. The libraries were pooled, and the concentrations were adjusted to 10 nM. The pooled libraries were denatured and neutralized by further dilution. Subsequently, the libraries were diluted to 1.8 pM and subjected to an NGS run on a NextSeq 500 System (Illumina, San Diego, CA, USA) with NextSeq500/550 v2.5 (75 Cycles) Kits (Illumina). Sequencing was performed using paired-end reads of 36 bases. After the sequencing run, FASTQ files were exported, and basic information of the NGS run data was checked for quality control using CLC Genomics Workbench 24.0 software (QIAGEN). The results, the PHRED-score as a quality score of the reads over 20 was confirmed for 99.8% of all reads, indicating successful data acquisition in the NGS run.

Supplementary Data S4. Bioinformatic analysis methods

Bioinformatic analysis was performed using CLC Genomics Workbench software, which proposed statistical analysis, obtained quantification values, and created heat maps, followed by mapping to the reference sequence of GCRm39 with GTF (general transfer format) file. Second, the data were analyzed using Microsoft Excel (Office 2019, Microsoft) to extract the genes of interest. Volcano plots were generated to detect significant differences in gene expression between the AR and AREx groups. Genes of interest were profiled using the RNA-seq data controller, highlighting expression features (RNA-seq chef)[14], a web-based integrated transcriptome analysis platform. For a detailed analysis, DEseq2 was used as the DEG analysis method and the Benjamini-Hochberg method was used as the false discovery rate (FDR) method. The trimmed mean M values (TMM) and transcripts per million (TPM) were normalized to the total read counts of the extracted genes of interest in the AR and AREx groups. The TMM-normalized values were used for gene set enrichment analysis (GSEA). GSEA was performed for all genes detected in each dataset using the GSEA web tool provided by the Broad Institute website (https://www.gsea-msigdb.org/gsea/index.jsp). GSEA was performed by selecting the Hallmark gene set from the Mouse Collection (MSigDB) and setting the AR group as 'Class A' and the AREx group as 'Class B'. The gene set database used in the degree of enrichment was quantified as an enrichment score (ES). A normalized enrichment score (NES) was calculated by normalizing the ES according to the size of the gene set. Data sets were compared using the NES: an NES greater than 2.0, and a small FDR indicated significant and strong enrichment. For the comparison of gene expression, TPM-normalized values were used between groups. The mouse immune cells were calculated by the single-sample Gene Set Enrichment Analysis (ssGSEA) method, as described previously with the gene set based on the R package of "ImmuCellAI-mouse".

Supplementary Data S5. Immunohistochemical staining

The sections were washed three times for 5 minutes each with PBS (pH, 7.4). For antigen activation, Proteinase K (Worthington Biochemical Co., Lakewood, NJ, USA)/distilled water (0.2 mg/mL) was added dropwise onto the sections and incubated for 15 min. After washing with PBS (three times for 5 min each), endogenous peroxidase activity was blocked by BLOXALL blocking solution (Vector Laboratories, Newark, CA, USA) for 10 min. The tissues were incubated for 30 min with 5% normal goat serum after washing in PBS (3 times, 5 min each). The following primary antibodies were incubated overnight at 4 °C: anti-CD4 rabbit polyclonal antibody (1:150 dilution, DF16080, Affinity Biosciences, Cincinnati, OH, USA), anti-tumor necrosis factor alfa (TNFα) rabbit polyclonal antibody (1:150 dilution, AF7014, Affinity Biosciences), anti-interleukin-1 beta (IL1β) rabbit polyclonal antibody (1:150 dilution, AF5103, Affinity Biosciences), anti-interleukin-6 (IL6) rabbit polyclonal antibody (1:200 dilution, ab6672, Abcam, Cambridge, UK), anti-interleukin-4 (IL4) rabbit polyclonal antibody (1:150 dilution, AF5142, Affinity Biosciences), and interleukin-10 (IL10) rabbit polyclonal antibody (1:100 dilution, DF6894, Affinity Biosciences). After washing with PBS (three times for 5 min each), the biotinylated goat anti-rabbit IgG (H+L) (1:100 dilution, BA-1000, Vector Laboratories) was used as a secondary antibody and incubated at room temperature for 30 min. After washing with PBS (three times for 5 min each), the avidin-biotin-peroxidase complex technique was performed at room temperature for 30 min using the Elite ABC Standard kit (PK-6100, Vector Laboratories). After washing with PBS three times for 5 min each, the sections were colored brown using a Peroxidase Stain DAB Kit (Brown Stain, NACALAI TESQUE, Kyoto, Japan). The nuclei were counterstained with Mayer's haematoxylins.

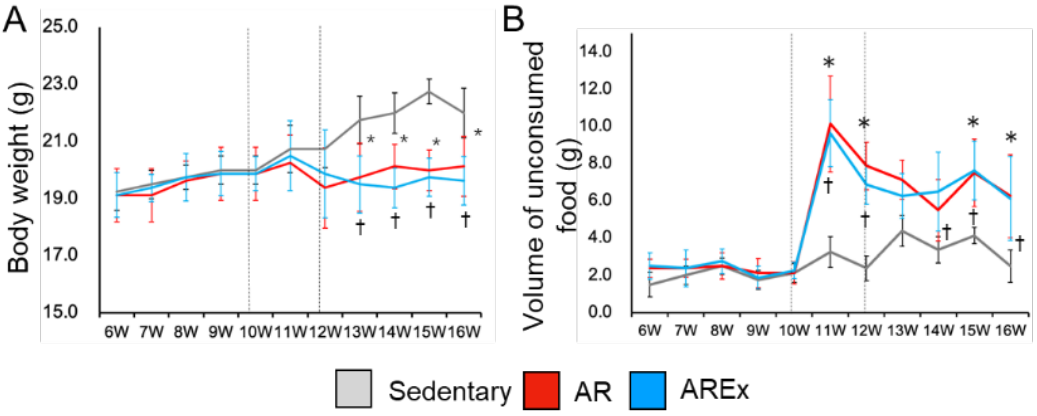


Supplementary Figure 1. Trends in body weight and food intake.

(A) Trends in body weight measured weekly from the start of rearing to the day of sacrifice in each mouse. (B) Trends in the volume of unconsumed food measured weekly from the start of rearing until the day of sacrifice in each mouse. Grey indicates the Sedentary group, red indicates the AR group, and blue indicates the AREx group. * ; Sedentary vs AR: $p<0.05$, †; Sedentary vs AREx: $p<0.05$, ‡; AR vs AREx: $p<0.05$.

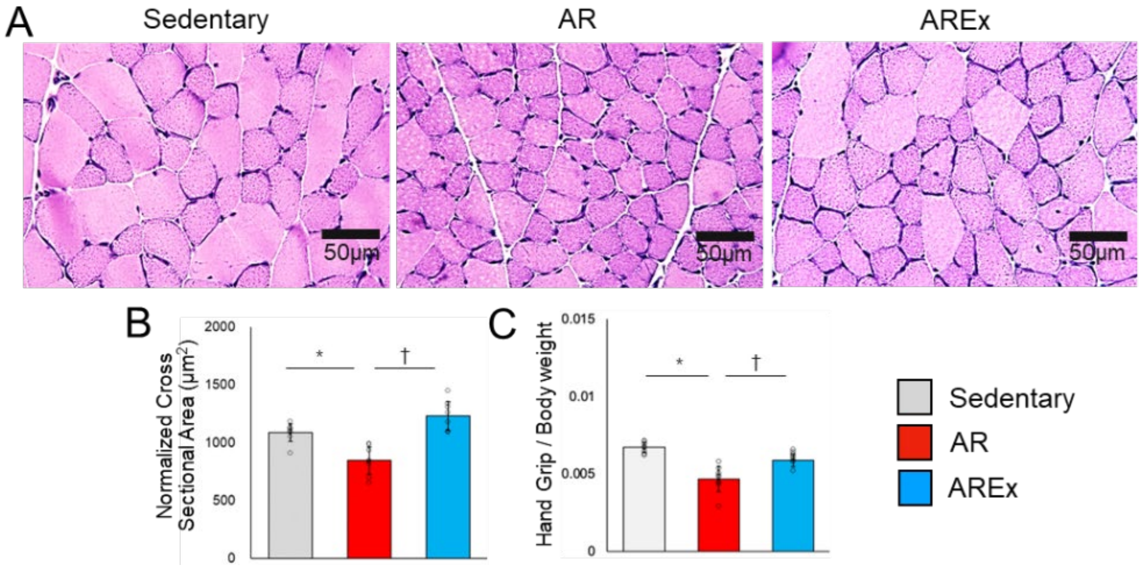


Supplementary Figure 2. Results of comparison of muscle transverse sectional area and handgrip strength.

(A) The histological images of the gastrocnemius muscle transverse sections with HE staining are shown. (B) Comparison of the average cross-sectional area of the gastrocnemius muscle. (C) Comparison of calculated handgrip strength normalized by body weight. Grey indicates the Sedentary group, red indicates the AR group, and blue indicates the AREx group. ＊; Sedentary vs AR: $p<0.05$, †; Sedentary vs AREx: $p<0.05$, ‡; AR vs AREx: $p<0.05$.

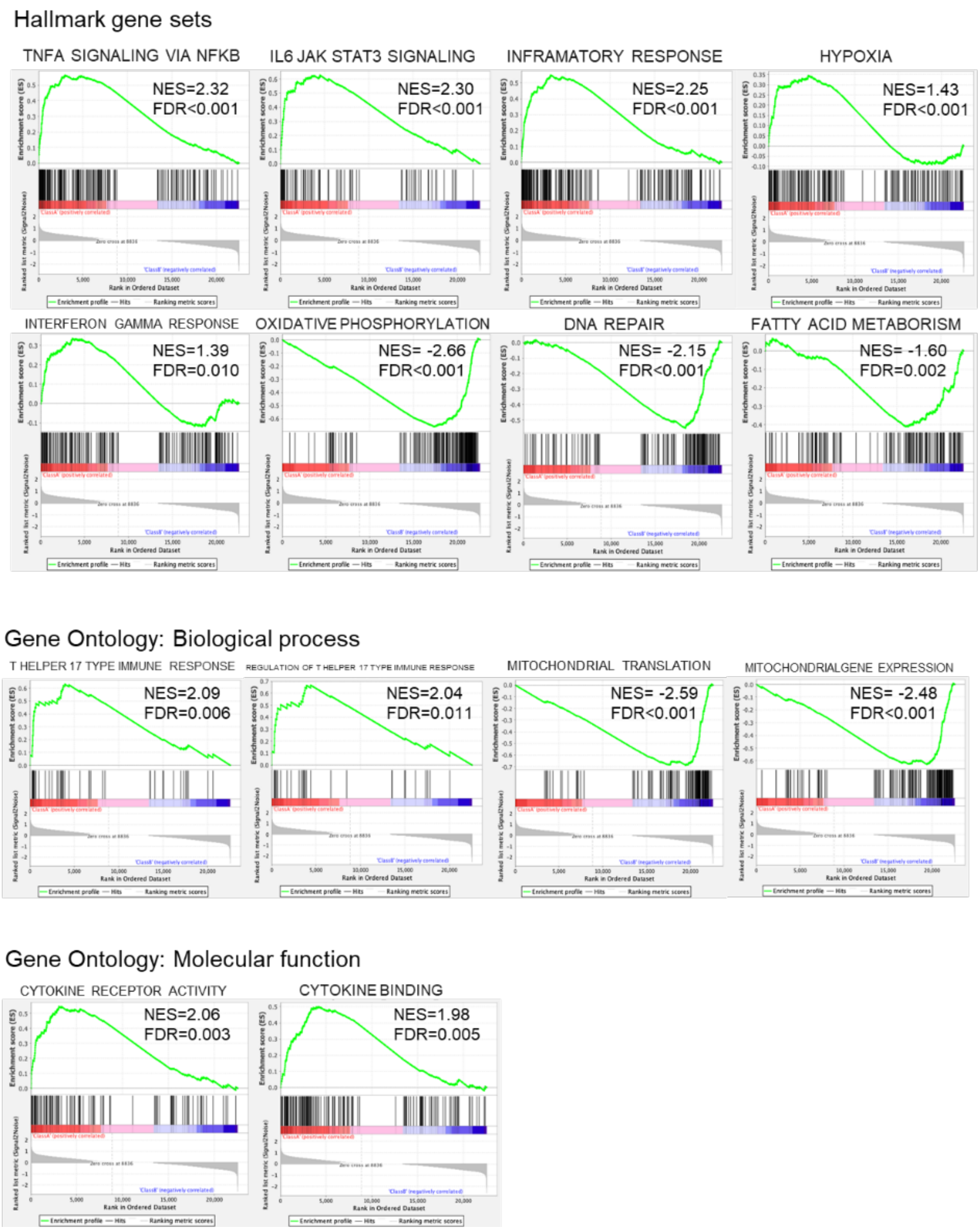


Supplementary Figure 3. GSEA results

TMM-normalized data from the total count values obtained from the bulk mRNA sequence were used. Some of the representative gene sets that ranked high in GSEA for the AR and AREx two-group comparisons were the hallmark gene set (Gene Ontology: biological process, molecular function). Red indicates the AR group as Class A, and blue indicates the AREx group as Class B. Normalized enrichment score (NES) and false discovery rate (FDR) values are shown.

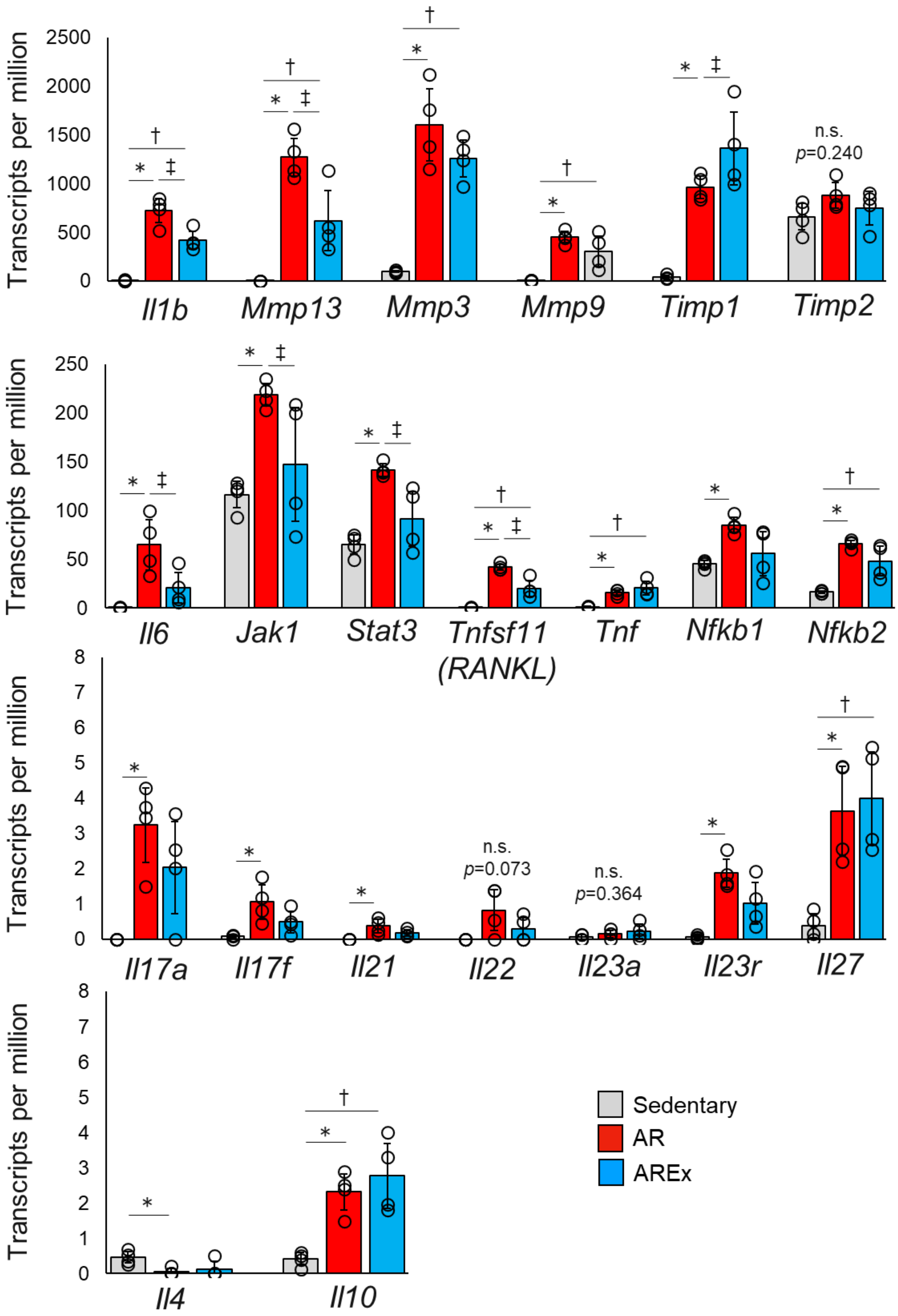


Supplementary Figure 4. TPM comparison for each gene.

TPM-normalized values obtained from the bulk mRNA sequence were used to compare specific gene variations between groups. Representative gene expression comparison results are presented. Grey indicates the Sedentary group, red indicates the AR group, and blue indicates the AREx group. ＊; Sedentary vs AR: $p<0.05$, †; Sedentary vs AREx: $p<0.05$, ‡; AR vs AREx: $p<0.05$.

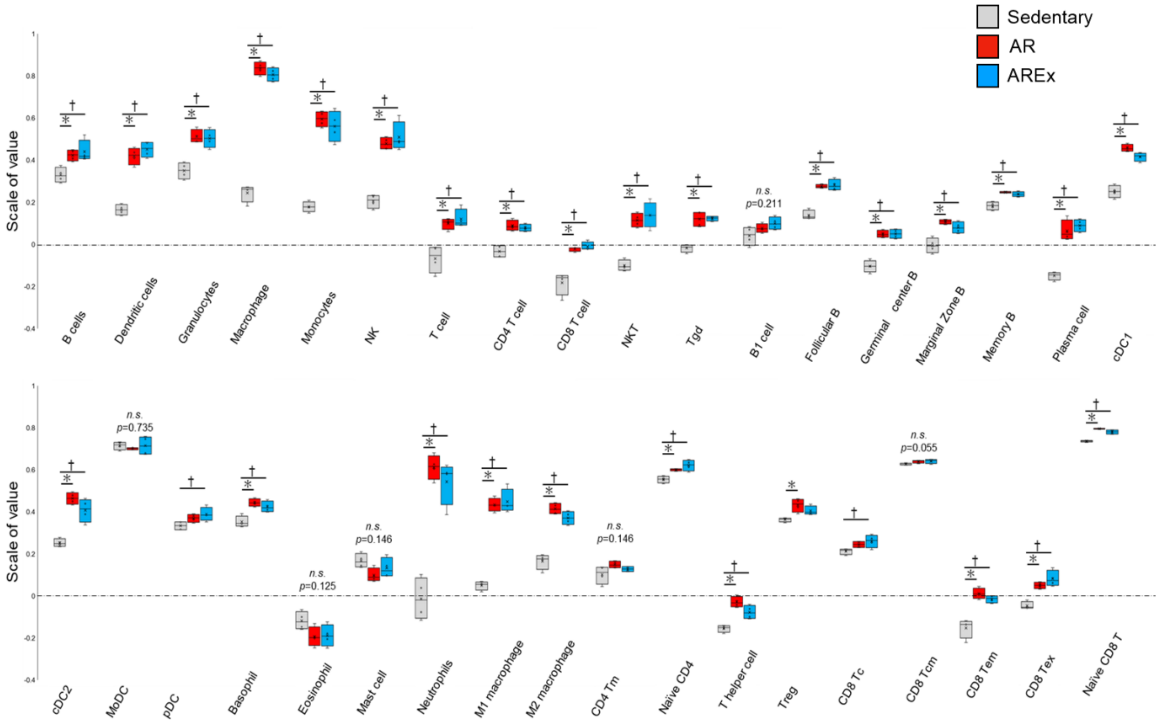


Supplementary Figure 5. Comparison of immune cell variability

For mouse immune cell variation, the single-sample Gene Set Enrichment Analysis (ssGSEA) method was used to calculate gene sets based on the R package ImmuCellAI-mouse against the values obtained from the bulk mRNA sequence. Grey indicates the Sedentary group, red indicates the AR group, and blue indicates the AREx group. ＊; Sedentary vs AR: $p<0.05$, †; Sedentary vs AREx: $p<0.05$, ‡; AR vs AREx: $p<0.05$.
